# Reversible and effective cell cycle synchronization method for studying stage-specific investigations

**DOI:** 10.1101/2024.09.02.610832

**Authors:** Yu-Lin Chen, Syon Reddy, Aussie Suzuki

## Abstract

The cell cycle is a crucial process for cell proliferation, differentiation, and development. Numerous genes and proteins play pivotal roles at specific cell cycle stages to regulate these events precisely. Studying the stage-specific functions of the cell cycle requires accumulating cell populations at the desired cell cycle stage. Cell synchronization, achieved through the use of cell cycle kinase and protein inhibitors, is often employed for this purpose. However, suboptimal concentrations of these inhibitors can result in reduced efficiency, irreversibility, and undesirable cell cycle defects. In this study, we have optimized effective and reversible techniques to synchronize the cell cycle at each stage in human RPE1 cells, utilizing both fixed high-precision cell cycle identification methods and high-temporal live-cell imaging. These reproducible synchronization methods are invaluable for investigating the regulatory mechanisms specific to each cell cycle stage.

## Introduction

Cell cycle is precisely regulated by a variety of kinases and proteins, with checkpoint mechanisms overseeing each stage to ensure proper cell cycle progression (Harper & Brooks, 2005; Schafer, 1998; Vermeulen *et al*, 2003). Disruption of this regulatory system can result in cancer and developmental diseases (Matthews *et al*, 2022). The reproductive cell cycle includes four major stages: G1, S, G2, and M phases, each with distinct functions. During the G1 phase, cells express proteins necessary for DNA synthesis, preparing for entry into the S phase. Cyclin D, in conjunction with Cdk4/6, plays a critical role in this process. The Cyclin D-Cdk4/6 complex phosphorylates the retinoblastoma protein (Rb), facilitating the release of Rb from E2F, an essential transcription factor (Harper & Brooks, 2005; Narasimha *et al*, 2014; Schafer, 1998; Vermeulen *et al*., 2003). This promotes E2F-dependent gene expression, including that of Cyclin E and Cyclin A, leading to the S phase entry. During the S phase, DNA polymerases orchestrate DNA replication. Cyclin E-Cdk2 promotes the transcription of histones, which are required for forming nucleosomes upon DNA synthesis (Armstrong *et al*, 2023; Harper & Brooks, 2005; Schafer, 1998; Vermeulen *et al*., 2003). After completing DNA replication, cells enter the G2 phase. The G2/M transition requires the activation of Cyclin B-Cdk1, and proper mitotic progression necessitates the degradation of Cyclin B (Harper & Brooks, 2005; Schafer, 1998; Vermeulen *et al*., 2003). The M phase, known as mitosis, includes five sub-stages: prophase, prometaphase, metaphase, anaphase, and telophase (Iemura *et al*, 2021).

Accumulating a cell population at the desired cell cycle stage is crucial for studying and identifying stage-specific gene/protein functions and interactions. One primary method for achieving this is fluorescence-activated cell sorting (FACS). FACS can sort cells based on specific cell cycle markers or DNA content in both live and fixed cells (Juan *et al*, 2002; Van Rechem *et al*, 2021). However, this technique requires specialized FACS equipment and a large number of cells, particularly when targeting low-abundance cell cycle stages, such as mitotic cells, in asynchronous populations (Whetstine & Van Rechem, 2022). Moreover, FACS often struggles to distinguish between the G2 and M phases and to identify detailed sub-stages within other cell cycle stages. Another widely used method involves cell cycle kinase and protein inhibitors (Banfalvi, 2011; Hadfield *et al*, 2022a; Wang, 2022). For example, Cdk4/6 inhibitors are extensively used in both basic research and clinical therapy for breast cancer, effectively arresting cells in the G1 phase (Wang *et al*, 2024). DNA polymerase inhibitors and DNA damage agents can arrest cells in the S phase, while Cdk1 inhibitors can halt cells in the G2 phase. Microtubule inhibitors are commonly used to synchronize cells in mitosis (Ligasova & Koberna, 2021). Although these cell cycle inhibitors are effective and user-friendly, it is crucial to use optimal concentrations and treatment durations. Using concentrations lower than optimal can lead to slower cell cycle progression with unintended defects, while higher concentrations can cause irreversible effects on the cell cycle. Both scenarios can potentially produce artificial results in experiments.

In this study, we carefully evaluate the effectiveness of widely used inhibitors for cell cycle synchronization at each stage of the cell cycle (G1, S, G2, and M phases). These synchronization protocols were specifically optimized for the hTERT-immortalized retinal pigment epithelial cell line (RPE1), a widely used, non-transformed human epithelial cell line in diverse research fields. By integrating a recently developed immunofluorescence (IF)-based cell cycle identification method (Chen *et al*, 2024) with high-temporal resolution live-cell imaging, we provide a comprehensive analysis of the impact of cell cycle arrest induced by major cell cycle inhibitors and their reversibility. The optimized cell synchronization techniques and thorough evaluation presented this study will be invaluable for investigating stage-specific regulatory mechanisms within the cell cycle.

## Results

### Cell cycle synchronization in G1 phase

We initially determined the detailed distribution of cell cycle phases in asynchronous RPE1 cells, which served as the standard in this study, using a recently developed high-precision, immunofluorescence-based cell cycle identification method (Chen *et al*., 2024) (**Supplementary Fig. 1a-b**). Cells were fixed and stained during the logarithmic growth phase (see **Methods**). An advantage of the use of IF-based cell cycle identification method allows us to determine detailed substages in cell cycle: G1, early S, late S, early G2, late G2, and each stage of mitosis, with a single cell resolution and accuracy. Our results revealed that approximately 50% of the cells were in the G1 phase, 20% in the early S phase, 10% in the late S phase, 11% in the early G2 phase, 4% in the late G2 phase, and 5% in mitosis (**Supplementary Fig. 1b**), aligning with previous results (Chen *et al*., 2024; Lau *et al*, 2009; McKinley & Cheeseman, 2017; Pei *et al*, 2022).

Effective and reversible cell cycle synchronization is crucial for studying protein functions associated with the cell cycle. This synchronization is typically achieved using chemical inhibitors that target kinase activities or essential proteins required for cell cycle progression (Mills *et al*, 2017; Wang, 2022). Cyclin-D, in conjunction with Cdk4/6, plays a pivotal role in regulating the G1 phase of cell cycle progression. The Cyclin-D-Cdk4/6 complex drives cell cycle progression by phosphorylating the Rb, thereby releasing the E2F transcription factor (Fassl *et al*, 2022). Previous research has demonstrated that Cdk4/6 inhibitors can induce G1 phase arrest in a wide variety of cells (Jost *et al*, 2021; Knudsen *et al*, 2020; Pennycook & Barr, 2021; Trotter & Hagan, 2020). Consequently, we investigated the detailed effects of Palbociclib, a highly selective Cdk4/6 inhibitor, on G1 cell cycle arrest (Liu *et al*, 2018). Prior studies have indicated that cells exposed to elevated concentrations of Palbociclib fail to resume cell cycle progression after washout (Trotter & Hagan, 2020). Therefore, we tested five concentrations of Palbociclib: 1, 0.5, 0.25, 0.1, or 0.05 µM. After treating cells with these concentrations of Palbociclib for 24 hours, they were subsequently subjected to the immunofluorescence-based cell cycle measurements (**Fig. 1a**). Our findings revealed that almost 100% of the cells treated with Palbociclib were arrested in G1 phase across a range of concentrations from 0.1 to 1 µM (**Fig. 1b**). However, when treated with 0.05 µM of Palbociclib, over 25% of the cells entered S phase, suggesting that this concentration is insufficient to fully arrest cells in G1 phase. We next investigated whether cells treated with Palbociclib could resume cell cycle progression following a washout. For this purpose, cells treated with Palbociclib for 24 hours were subjected to a washout process and then exposed to STLC, an Eg5 inhibitor known to induce mitotic arrest, for an additional 18 hours. After this period, cells were fixed and assessed the cell cycle distribution (**Fig. 1c and Supplementary Fig. 2a**). Our findings revealed that cells treated with concentrations ranging from 0.05 to 0.5 µM of Palbociclib demonstrated a 50-60% incidence of the S phase and up to 20% of cells in mitosis, suggesting that these concentrations enable the resumption of cell cycle progression. However, approximately 30% of cells treated with these concentrations remained arrested in the G1 phase. In contrast, treatment with 1 µM Palbociclib resulted in a significantly higher proportion of cells in the G1 phase (approximately 55%), indicating an impaired ability to restart cell cycle progression at this concentration. To corroborate these results, we employed live-cell imaging using RPE1 H2B-EGFP cells immediately following the Palbociclib washout (**Fig. 1d**). In alignment with the immunofluorescence quantifications, cells exposed to Palbociclib at concentrations ranging from 0.1 to 0.5 µM entered mitosis approximately 12 to 15 hours post-washout (**Fig. 1d, arrows**). Conversely, cells treated with 0.05 µM Palbociclib exhibited mitotic cells as early as 9 hours after washout, while those treated with 1 µM rarely showed signs of mitosis. To summarize, our study suggests that Palbociclib concentrations ranging 0.1 µM to 0.5 µM, which effectively induce G1 phase arrest, allow cells to resume cell cycle progression following washout in RPE1 cells.

**Figure 1:**
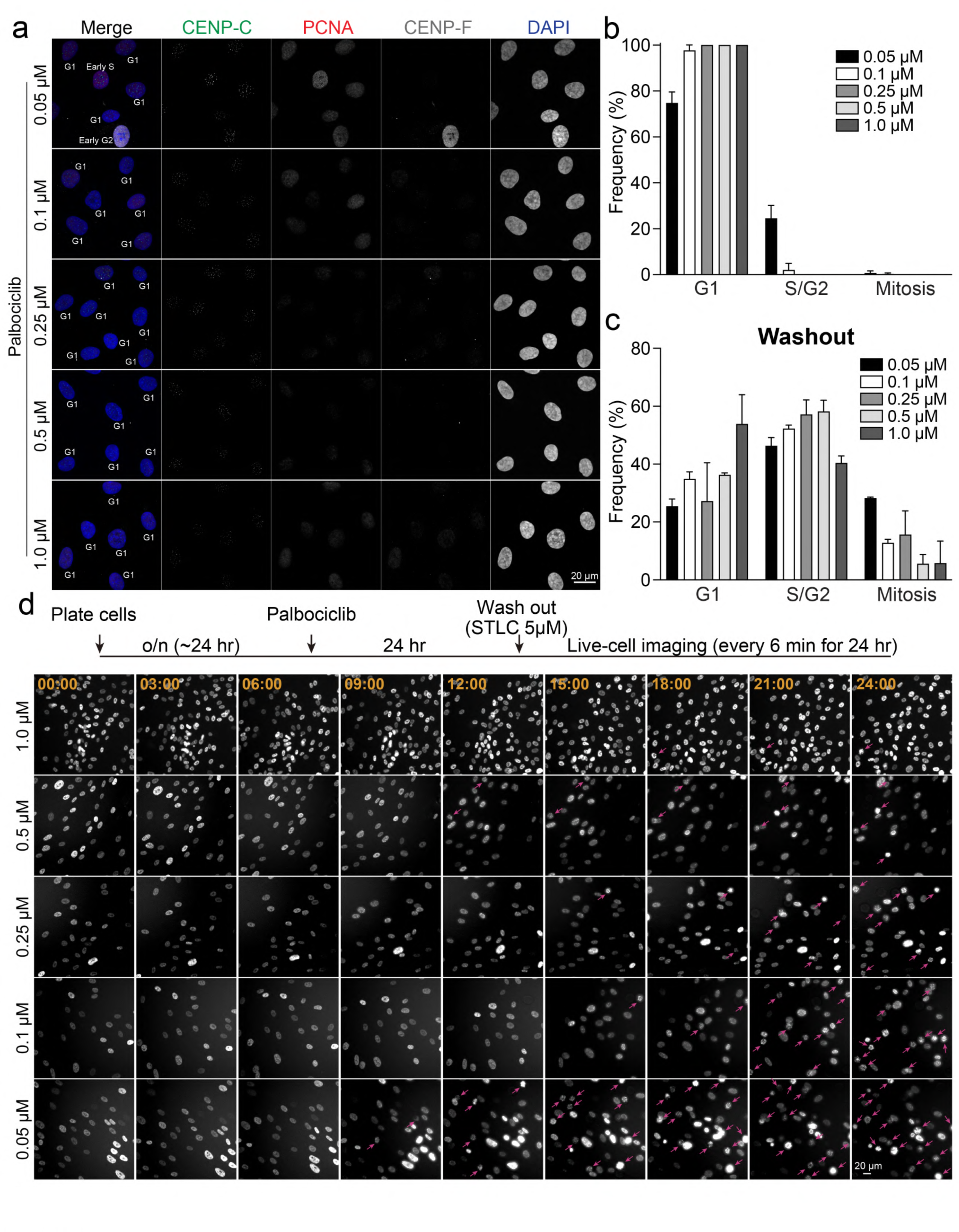
**G1 phase synchronization and release by Palbociclib** (a) Representative immunofluorescence images of RPE1 cells treated with Palbociclib conditions (0.05, 0.1, 0.25, 0.5, or 1 µM for 24 hours), labeled with antibodies for CENP-C, PCNA, and CENP-F. (b) Proportion of RPE1 cells in G1, S/G2, or M phase in condition (a). From left to right, n = 416, 408, 383, 417, 487 (from two replicates). (c) Proportion of RPE1 cells in G1, S/G2, or M phase, analyzed at 18 hours following the washout of Palbociclib. From left to right, n = 369, 317, 336, 355, 393 (from two replicates). (d) Schematic timeline of live-cell imaging sequence (top). Representative live-cell imaging of H2B-GFP expressing RPE1 cells treated with Palbociclib (0.05, 0.1, 0.25, 0.5, or 1 µM for 24 hours) (Bottom). Palbociclib was washed out prior to imaging. Mitotic cells are indicated with pink arrows. Imaging was performed at least two independent replicates.

### Cell cycle synchronization in S phase

Aphidicolin, a tetracyclic diterpene antibiotic, specifically inhibits DNA polymerases, enzymes essential for DNA replication during the S phase (Ikegami *et al*, 1978; Krokan *et al*, 1981). The effect of Aphidicolin on cell cycle progression has been a subject of debate, with varying studies presenting contradictory findings. Some research posits that Aphidicolin induces an arrest in the early S phase (Bhaud *et al*, 2000; Fragkos *et al*, 2019; Maeda *et al*, 2014; Mazouzi *et al*, 2016; Xu *et al*, 2011; Xu *et al*, 2001), whereas others suggest it causes cells to halt at the G1 phase, likely right on the cusp of the G1-S transition (Engstrom & Kmiec, 2008; Saintigny *et al*, 2001; Szczepanski *et al*, 2019; Yiangou *et al*, 2019).

To elucidate the precise impact of Aphidicolin on cell cycle progression, we conducted immunofluorescence-based cell cycle analysis using RPE1 cells. Our experiments involved a 24-hour treatment with Aphidicolin at concentrations of 2.5, 5, or 10 µg/ml. We found that approximately 90% of Aphidicolin-treated cells showed an absence of punctuated PCNA and CENP-F nuclear signals across all concentrations, indicating that Aphidicolin arrests RPE1 cells in G1 phase rather than S phase (**Fig. 2a-b and Supplementary Fig. 2c-d**). Consistent with these findings, live-cell imaging revealed that cells treated with Aphidicolin at concentrations of 2.5 or 5 µg/ml did not exhibit any mitotic entry after 9 hours of treatment, whereas control cells continued to enter mitosis within the 24-hour imaging period (**Supplementary Fig. 2b**). These results suggest that Aphidicolin effectively inhibits the initiation of DNA replication and arrests RPE1 cells in G1 phase.

**Figure 2:**
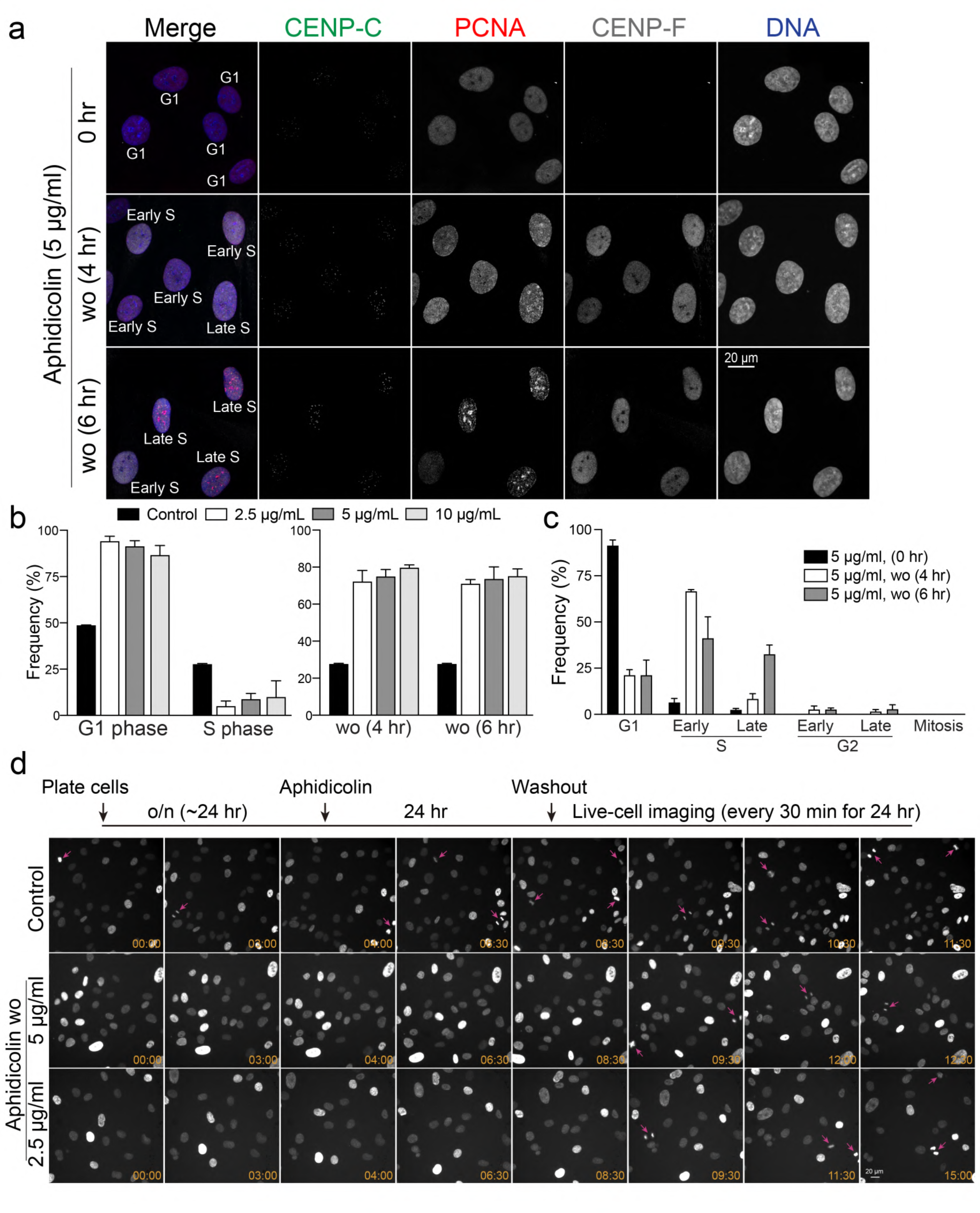
**S phase synchronization and release in RPE1 cells using Aphidicolin** (a) Representative immunofluorescence images of RPE1 cells treated with control or aphidicolin conditions, labeled with antibodies for CENP-C, PCNA, and CENP-F. Images captured before and at 4 or 6 hours post-aphidicolin washout. (b) Proportion of RPE1 cells in G1 or S phase, analyzed before (left) and at 4 or 6 hours (right) following the washout of aphidicolin (treated at concentrations of 2.5, 5, or 10 µg/ml for 24 hours). For left panel, from left to right, n = 424, 400, 370, 344 (from two replicates). For right panel, from left to right, n = 424, 379, 350, 341, 424, 401, 375, 413 (from two replicates). Data represented from two experimental replicates. (c) Proportion of cells in distinct cell cycle stages (G1, Early S, Late S, Early G2, Late G2, and Mitosis) before and after 4 or 6 hours post-aphidicolin washout (5 µg/ml). From left to right, n = 370, 350, 375 (from two replicates). (d) Schematic timeline of live-cell imaging sequence (top). Representative live-cell imaging of H2B-GFP expressing RPE1 cells treated with either DMSO (control) or aphidicolin (2.5 or 5 µg/ml for 24 hours), (bottom). Aphidicolin was washed out prior to imaging. Mitotic cells are indicated with pink arrows. Imaging was performed at least two independent replicates.

To achieve S phase synchronization, we aimed to determine the timing and conditions under which cells could enter the S phase following the removal of Aphidicolin. For this purpose, we incubated cells with Aphidicolin at concentrations of 2.5, 5, or 10 µg/ml for 24 hours, and subsequently fixed and stained the cells at 4 or 6 hours after removing Aphidicolin. Our results showed that approximately 80% of the cells entered the S phase at both 4 and 6 hours post-Aphidicolin removal across all tested concentrations (**Fig. 2a-c**). Specifically, at 4 hours post-Aphidicolin washout at a concentration of 5 µg/ml, approximately 67% of cells were in early S phase and 10% were in late S phase (**Fig. 2c**). This shifted to 49% in early S phase and 29% in late S phase by 6 hours (**Fig. 2c**). Similar trends were observed in cells treated with 2.5 or 10 µg/ml at 4 or 6 hours after removal of Aphidicolin (**Supplementary Fig. 2d**). These observations demonstrate a dynamic recovery, with about 80% of RPE1 cells successfully progressing to the S phase within 4 to 6 hours after a 24-hour exposure to Aphidicolin at concentrations ranging from 2.5 to 10 µg/ml. To further validate these results, we conducted live-cell imaging following Aphidicolin washout (**Fig. 2d**). Mitotic cells appeared only 9 hours after Aphidicolin washout, whereas control cells continued to exhibit mitotic cells during live imaging (**Fig. 2d, arrows**). This corresponds to the results obtained from the fixed immunofluorescence-based cell cycle analysis (**Fig. 2a-c**). In summary, our study not only dissects the cell cycle arrest induced by Aphidicolin but also highlights its capability for effective S phase synchronization. Aphidicolin removal is effective for studies focusing on early S phase within 4 hours, and on late S phase after more than 6 hours.

### Cell cycle synchronization in G2 phase

The Cyclin B-Cdk1 complex orchestrates both mitotic entry and exit. To initiate mitosis, Cyclin B-Cdk1 must be activated by Cdc25 phosphatase, which dephosphorylates Cdk1, converting it from its inactive to active form (Vassilev, 2006). Inhibition of Cdk1 prior to mitosis prevents mitotic entry (Lau *et al*, 2021). Supporting this, the small-molecule inhibitor of Cdk1, RO-3306, effectively arrests cells in G2 phase, as observed through flow cytometry (Johnson *et al*, 2021; Tanenbaum *et al*, 2015; Vassilev *et al*, 2006). We tested various concentrations of RO- 3306 in RPE1 cells to analyze the specific cell cycle stages arrested. Cells were incubated with 1, 3, 6, or 10 µM of RO-3306 for 24 hours, fixed, and then the cell cycle stages were determined using an immunofluorescence-based cell cycle identification method (**Fig. 3a**). We found that treatment with 3 and 6 µM RO-3306 efficiently accumulated cells in the G2 phase, with 60% and 58% of cells respectively, while only 12-13% of cells accumulated in G2 at 1 and 10 µM (**Fig. 3b**). Surprisingly, most cells treated with 10 µM RO-3306 were arrested in the G1 phase (**Fig. 3c**), indicating that a high concentration of RO-3306 may inhibit other Cdks in addition to its primary target, Cdk1 (Jorda *et al*, 2018). In RPE1 cells, 1 µM of RO-3306 was insufficient to arrest cells in the G2 phase (**Fig. 3c**). Treatment with 3 µM RO-3306 resulted in nearly equal populations of early and late G2 phase cells (28% and 32%, respectively), whereas 6 µM RO- 3306 predominantly arrested cells in early G2 phase (**Fig. 3c**). Notably, we observed a subset of interphase cells exhibiting bubbled nuclei specifically in the 3 µM RO-3306-treated groups (**Fig. 3d**). Next, we examined the mitotic index after RO-3306 washout. We quantified mitotic cells at 2 hours post-washout in STLC-contained growth medium. Cells treated with 3 µM RO-3306 exhibited ∼30% mitotic cells at 2 hours post-washout. Interestingly, only ∼8% of cells treated with 6 µM RO-3306 entered mitosis within 2 hours of washout, and no mitotic cells were observed after washout in cells treated with 10 µM RO-3306 (**Fig. 3e**), suggesting that cells cannot efficiently recover at these concentrations.

**Figure 3:**
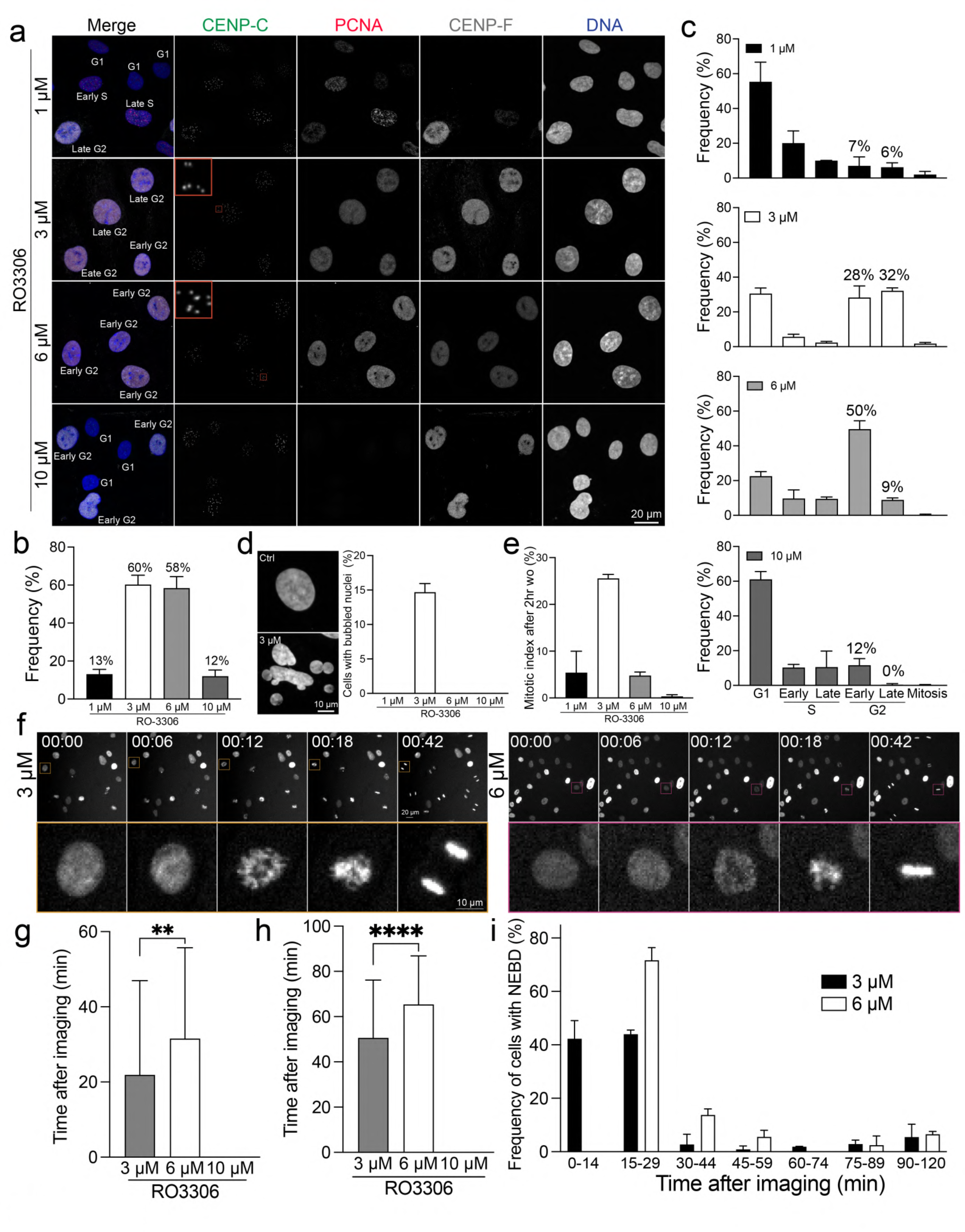
**G2 phase synchronization and release in RPE1 cells using RO-3306** (a) Representative immunofluorescence images of RPE1 cells under control conditions or treated with RO-3306 (1, 3, 6, or 10 µM for 24 hours), stained with antibodies for CENP-C, PCNA, and CENP-F. (b) Percentage of cells in G2 phase in condition (a). From left to right, n = 404, 362, 318, 388 (from two replicates). (c) Proportion of cells in each stage of cell cycle in condition (a). (d) Representative DNA images and percentage of cells with bubbled nucleus. From left to right, n = 404, 362, 318, 388 (from two replicates). (e) Mitotic index at 2 hours after RO-3306 washout in growth media containing STLC. From left to right, n = 739, 483, 508, 601 (from two replicates). (f) Representative live-cell imaging of H2B-GFP expressing RPE1 cells treated with either DMSO (control) or varying concentrations of RO-3306 (3 or 6 µM for 24 hours). RO-3306 was washed out prior to imaging. (g) Average time to nuclear envelope breakdown (NEBD) post-imaging initiation, in cells treated with either DMSO (control) or RO-3306 at concentrations of 3, 6, and 10 µM for 24 hours. The RO-3306 treatment was washed out before imaging commenced. n = 107 and 93 (from left to right, three replicates). 10 µM of RO3306 washout did not show any mitotic cells in two independent replicates. (h) Average time to anaphase onset in cells from condition (g). n = 107 and 90 (from left to right, three replicates). 10 µM of RO3306 washout did not show any mitotic cells in two independent replicates. (i) The proportion of cells that enter NEBD after the start of imaging for the same treatments of (g). n = 107 and 93 (from left to right, three replicates).

To further validate our quantification results obtained in fixed cell analysis, we performed live-cell imaging using H2B-GFP-expressing RPE1 cells immediately after treatment of 3 or 6 µM RO-3306 (**Supplementary Fig. 3a**). While control cells consistently exhibited mitotic progression during live-cell imaging, cells treated with 6 µM RO-3306 did not show any progress to mitosis, indicating that 6 µM of RO-3306 effectively inhibits mitotic entry. Although mitotic index was significantly reduced in cells treated with 3 µM RO-3306, the subset of cells that entered mitosis experienced a slight but significant delay in mitotic duration and nuclear bubbling (**Supplementary Fig. 3a (arrow) and 3b**), consistent with observations in fixed-cell analysis (**Fig. 3d**). These results demonstrate that 3 µM and higher concentration of RO-3306 efficiently arrests most cells in G2 phase, but a subset of these G2 phase cells can enter mitosis. These mitotic cells displayed significant errors in both mitotic progression and anaphase, resulting in nuclear bubbling (**Supplementary Fig. 3c**) (Voets *et al*, 2015).

Next, we examined recovery after RO-3306 washout using live-cell imaging (**Supplementary Fig. 3d**). Both 3 µM and 6 µM RO-3306-treated cells exhibited NEBD and anaphase onset approximately 20-30 minutes and 50-70 minutes, respectively, after RO-3306 washout. In contrast, no mitotic cells were observed in the presence of 10 µM RO-3306 (**Fig. 3e, g, and h, Supplementary Fig. 3d**). After washout, cells treated with 3 µM RO-3306 entered mitosis significantly faster than those treated with 6 µM (**Fig. 3f-i**). Collectively, RO-3306 at concentrations between 3 to 6 µM effectively accumulate cells in G2 phase, and 3 µM RO-3306 provides better recovery after washout. Higher concentrations of RO-3306 (10 µM in RPE1 cells) fail to synchronize RPE1 cells in G2 phase and prevent, at least efficient, recovery to a normal cell cycle progression even after RO-3306 removal.

### Cell cycle synchronization in Prometaphase

Microtubule depolymerizers, including Nocodazole and Colcemid, have traditionally been used for mitotic synchronization due to their ability to effectively disrupt spindle formation and prevent chromosome segregation (Florian & Mitchison, 2016; Hadfield *et al*, 2022b; Surani *et al*, 2021). However, despite their reversible nature, cells treated with these drugs and subsequently washed exhibit a marked increase in severe mitotic defects due to the lack of microtubule dynamicity (Cavazza *et al*, 2016; Worrall *et al*, 2018). Due to these limitations, our study employed STLC, a potent Eg5 inhibitor, as an alternative agent to arrest cells in mitosis (Florian & Mitchison, 2016; Hadfield *et al*., 2022b). Following NEBD, chromosomes undergo dynamic interactions with microtubules during prometaphase, including the capture of kinetochores and the establishment of bipolar spindles required for metaphase plate formation. While high concentrations of traditional microtubule depolymerizers obliterate microtubules, Eg5 inhibitors do not prevent microtubule assembly at kinetochores. Instead, it impedes centrosome separation necessary for bipolar spindle formation, resulting in prometaphase arrest while maintaining kinetochore-microtubule interactions (Skoufias *et al*, 2006). Consequently, removing Eg5 inhibitors is thought to facilitate a more effective recovery than treatment with microtubule depolymerizers (Bakhoum *et al*, 2009).

In our study, we treated cells with 2, 5, or 10 µM STLC for 24 hours and assessed the mitotic index. The results showed that 5 and 10 µM concentrations achieved approximately 60% synchronization efficiency, whereas 2 µM STLC treatment exhibited nearly equivalent synchronization efficiency as untreated control (**Fig. 4a and 4b**). As expect, in the presence of 5 and 10 µM STLC, almost 100% of the mitotic cells were arrested in prometaphase and exhibited monopolar spindles (**Fig. 4c and 4d**). These results confirm the efficiency of 5 and 10 µM STLC in synchronizing cells at prometaphase. For applications requiring a higher purity of prometaphase populations, we recommend using a mitotic shake-off technique (Zwanenburg, 1983) following STLC synchronization, which yielded nearly 100% pure prometaphase population (**Fig. 4a and 4e**). We validated the immunofluorescence-based quantification of STLC synchronization by live-cell imaging. RPE1 cells treated with 5 or 10 µM STLC demonstrated a gradual and efficient accumulation in prometaphase, with ∼80% of cells arrested in this stage after 24 hours (**Fig. 4f (arrows), 4g, and Supplementary Fig. 3e**). Nearly 100% of these prometaphase cells formed monopolar spindles due to Eg5 inhibition (**Fig. 4h**). In contrast, most cells treated with 1 µM of STLC could proceed through division (**Fig. 4g and Supplementary Fig. 3e**). Importantly, there was no significant increase in apoptotic cell death among cells treated with any concentration of STLC compared to the control during 24 hours of live imaging (**Supplementary Fig. 3e**). These observations are in alignment with the results obtained from immunofluorescence-based quantifications, which showed that treatment with 5 and 10 µM of STLC effectively arrests cells in prometaphase.

**Figure. 4:**
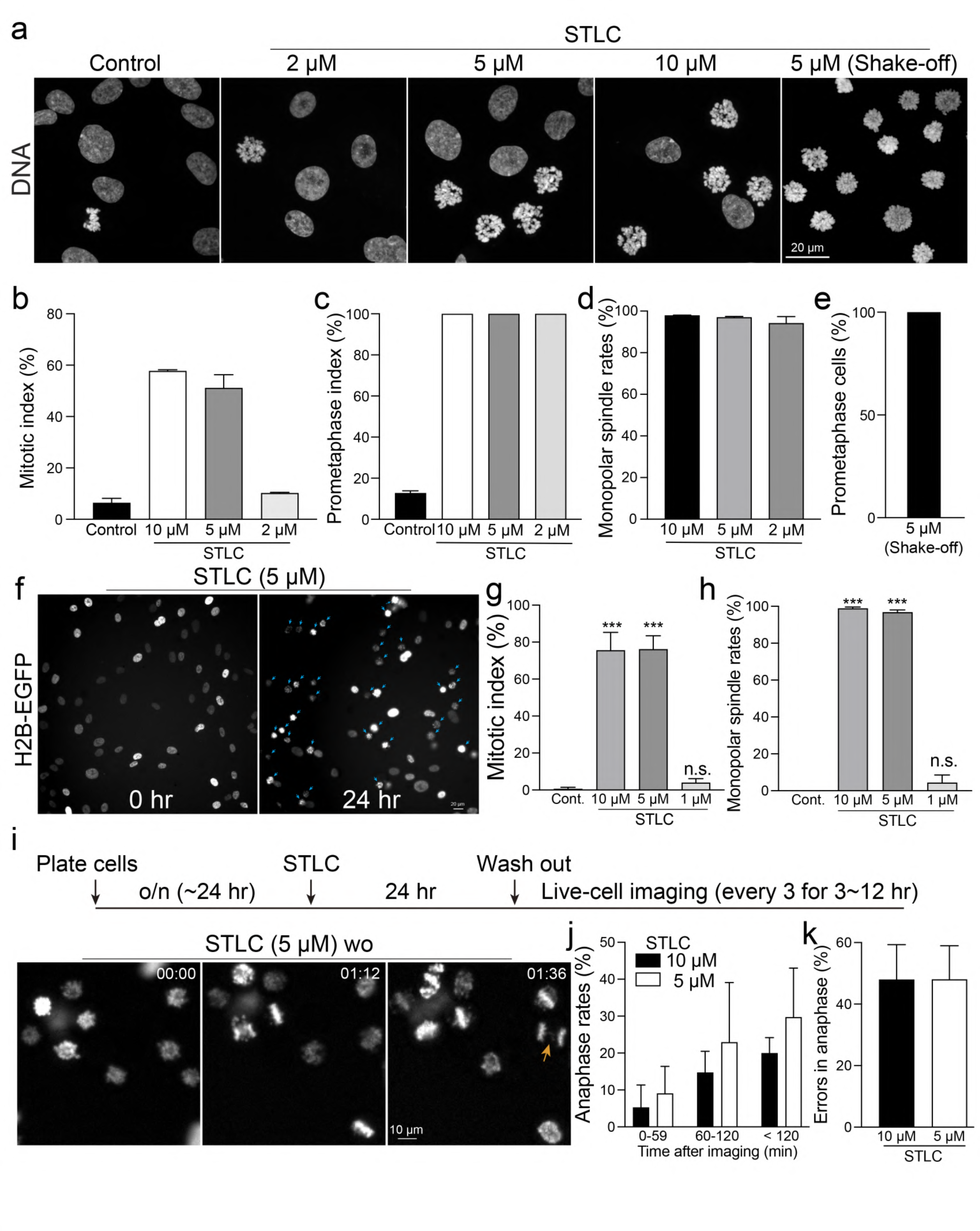
**Prometaphase synchronization and release in RPE1 cells using STLC** (a) Representative confocal images of DNA in RPE1 cells under control conditions, treated with STLC at concentrations of 2, 5, or 10 µM, and post-mitotic shake-off following treatment with 5 µM STLC. (b) Mitotic index of cells under control conditions compared to those treated with STLC (2, 5, or 10 µM). From left to right, n = 424, 452, 406, 789 (two replicates). (c) Prometaphase index corresponding to the treatments described in (b). From left to right, n = 341, 261, 208, 81 (two replicates). (d) Percentage of mitotic cells displaying a monopolar spindle after treatment with STLC at 2, 5, or 10 µM. From left to right, n = 193, 204, 65 (two replicates). (e) Mitotic index following mitotic shake-off in cells treated with 5 µM STLC. n = 503 (two replicates). (f) Representative live-cell imaging of H2B-GFP-expressing RPE1 cells treated with 5 µM STLC. (g) Mitotic index in live cells under control conditions and after treatment with STLC at concentrations of 1, 5, or 10 µM for 24 hours. (h) Proportion of mitotic cells with a monopolar spindle following the treatments outlined in (g). (i) Schematic timeline of live-cell imaging sequence (Top). Representative live-cell imaging of H2B-GFP expressing RPE1 cells treated with either DMSO (control) or 5 µM STLC for 24 hours, after which STLC was washed out (Below). A mitotic cell with lagging chromosomes is highlighted with an orange arrow. (j) Proportion of cells progressing to anaphase. (k) Percentage of anaphase cells exhibiting errors, including lagging chromosomes and chromosome bridges. n = 679 and 518 (from left to right (j and k), two replicates)

We next investigated whether mitotic cells arrested by STLC could exit mitosis after washout. For this experiment, RPE1 cells were incubated with STLC at concentrations of 5 µM or 10 µM for 24 hours. Following the washout, we immediately commenced high-temporal- resolution live-cell imaging (**Fig. 4i and Supplementary Fig. 3f**). We quantified the percentage of arrested cells that entered anaphase within 2 hours post-washout. Our results showed that approximately 20% and 30% of the cells arrested in prometaphase progressed to anaphase within 2 hours after washout of 5 µM or 10 µM STLC, respectively (**Fig. 4j**). Notably, only 10% of cells underwent anaphase within the first hour. Among these divided cells, about 50% exhibited errors during anaphase (**Fig. 4i (arrow)-k, and Supplementary Fig. 3f (arrow)**). These findings indicate that only a subset of STLC-arrested cells is able to enter anaphase immediately after the washout.

### Cell cycle synchronization in Metaphase, Anaphase, and Telophase

The transition from metaphase to anaphase necessitates the degradation of Cyclin B and Securin (Han & Li, 2014). This degradation activates Separase, allowing it to cleave the cohesion between sister chromatids and enabling their segregation. Consequently, proteasome inhibitors such as MG132 have been identified to effectively induce metaphase arrest (Daum *et al*, 2011; Santamaria *et al*, 2007; Tipton & Gorbsky, 2022). Previous studies have demonstrated that cells treated with MG132 maintain the metaphase plates, resulting in kinetochores experiencing heightened tension compared to those in normal metaphase (Wan *et al*, 2009). This increased tension is evidenced by the observed increases in the intra- and inter-kinetochore stretch. However, it is important to note that proteasome inhibitors lack specificity in mitotic processes, raising concerns about their potential to disrupt various cell cycle regulations inadvertently. To support this, unlike STLC, RPE1 cells treated with 10 µM MG132 for 24 hours did not show a significant increase in mitotic index (**Fig. 5a and 5b**). On the other hand, metaphase cells exposed to long-term MG132 treatment exhibited significant defects in chromosome alignment (**Fig. 5a and 5c**), likely due to cohesion fatigues (Daum *et al*., 2011). To further validate this observation, we performed live-cell imaging on cells treated with 10 µM MG132 (**Fig. 5d**). Although these cells established and maintained a metaphase plate for approximately 2 hours after NEBD, the spatial organization of chromosomes became disorganized thereafter, leading to misaligned chromosomes and apoptotic cell death. These results demonstrate that using MG132 alone is insufficient for synchronizing cells in metaphase, anaphase, and telophase. To enrich populations of metaphase cells, we utilized a combination approach involving RO-3306 for G2 cell synchronization followed by MG132 treatment (**Fig. 5e and Supplementary Fig. 4a**). As the majority of cells arrested by RO-3306 progress to NEBD within 1 to 2 hours, we investigated the effects of MG132 treatments for 1 or 2 hours on the synchronization efficacy of metaphase cells following RO-3306 washout. Our findings reveal that the combination of RO-3306 and MG132 effectively increases the population of metaphase cells (**Fig. 5e and Supplementary Fig. 4a**). Interestingly, approximately 40-50% of cells arrested in metaphase after 2 hours of MG132 treatment fail to initiate anaphase within 2 hours after MG132 washout (**Fig. 5f**). In contrast, nearly 100% of these cells subjected to 1-hour MG132 treatment enter anaphase. This phenotype is not rescued by reducing the concentration of MG132 to 5 µM, suggesting that MG132 treatment exceeding 1 hour or arresting cells in metaphase for longer than 1 hour impedes anaphase entry even after washout.

**Fig. 5:**
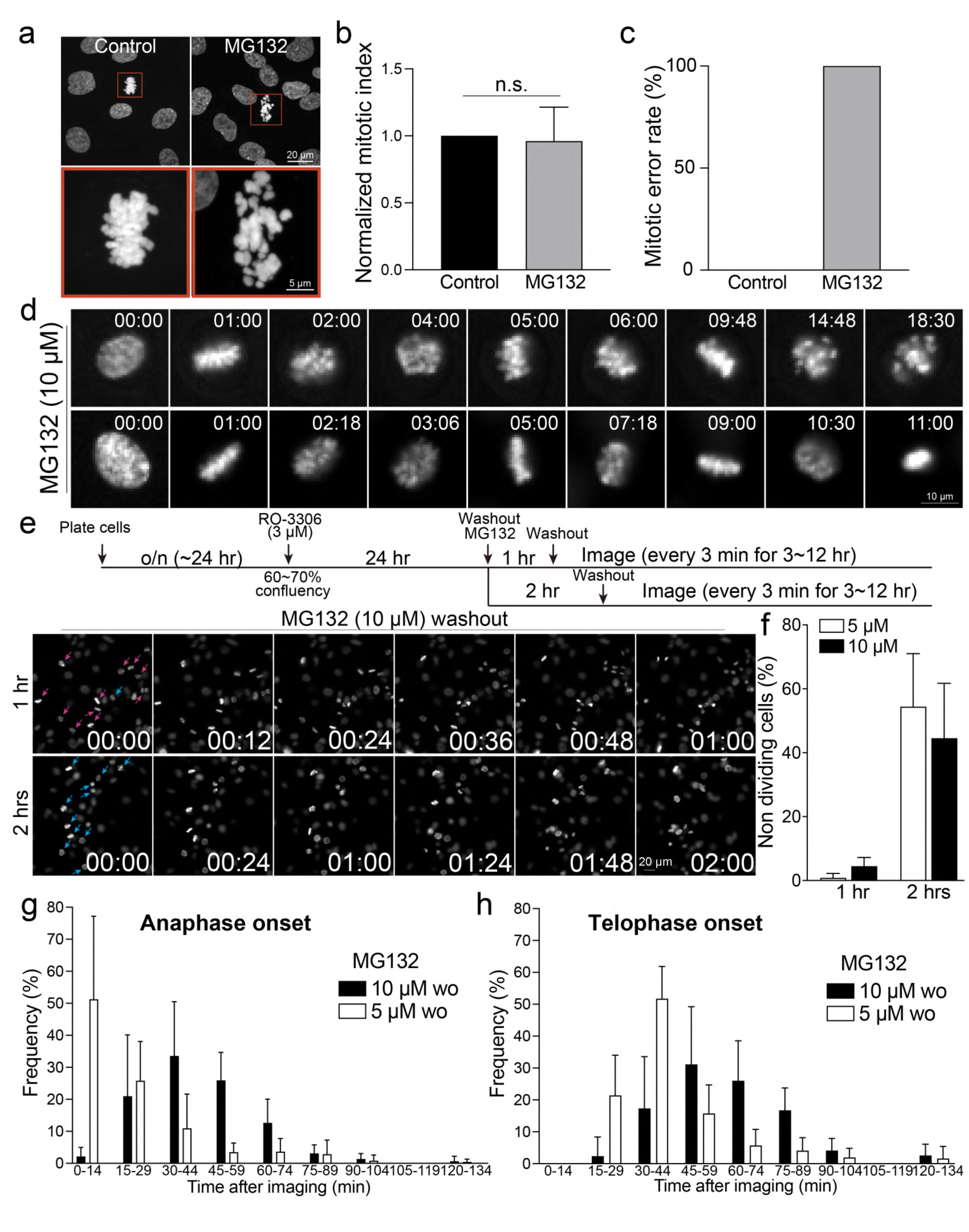
Metaphase, Anaphase, and Telophase synchronization using both RO-3306 and MG132. (a) Representative confocal images of DNA in RPE1 cells under control conditions or treated with MG132 (10 µM) for 24 hours. (b) Mitotic index of cells under control conditions compared to those treated with MG132 (10 µM) for 24 hours. From left to right, n = 619, 647 (two replicates). (c) Mitotic error rates in control or cells treated with MG132 (10 µM) for 24 hours. n = 12, from two replicates. (d) Representative live-cell imaging of H2B-GFP-expressing RPE1 cells treated with 10 µM MG132. (e) Schematic timeline of live-cell imaging sequence (Top). Representative live-cell imaging of H2B-GFP expressing RPE1 cells treated with10 µM MG132 for either 1 or 2 hours, after which MG132 was washed out (Bottom). Prior to the treatment of MG132, cells were incubated with RO-3306 for 24 hours. (f) Proportion of non-dividing mitotic cells following the treatments outlined in (e). n = 196, 237, 257, 164 (from left to right, two replicates). (g and h) Proportion of cells entering anaphase onset or telophase onset in condition (e). n = 237, 196 (from two replicates).

For anaphase cell synchronization, cells treated with 5 µM of MG132 for 1 hour exhibited anaphase onset immediately after washout, with approximately 80% of cells entering anaphase within 30 minutes after MG132 removal (**Fig. 5g and Supplementary Fig. 4b**). Conversely, cells treated with 10 µM of MG132 showed approximately 60% of cells entering anaphase within a range of 30 to 60 minutes after washout. The telophase population peaked between 30 and 60 minutes in cells treated with 5 µM MG132 and between 45 and 75 minutes in cells treated with 10 µM MG132 after washout (**Fig. 5h and Supplementary Fig. 4c**). About 50% of anaphase cells exhibited errors in both 5 and 10 µM MG132-treated cells for 2 hours, whereas approximately 16-30% of these cells exhibited errors after 1 hour of treatment (**Supplementary Fig. 4d**). Although no metaphase-arrested cells treated with 5 µM MG132 for 1 or 2 hours exhibited apoptotic cell death within 2 hours after washout, 2-5% of cells exhibited apoptotic cell death in cells treated with 10 µM MG132 for 1 and 2 hours, respectively (**Supplementary Fig. 4e**). Additionally, no anaphase cells were found in cells treated with 10 µM MG132 at the beginning of imaging, while cells treated with 5 µM MG132 for 1 hour occasionally entered mitosis upon imaging (**Supplementary Fig. 4f**). Collectively, the combination of RO-3306 G2 cell cycle synchronization and a 1-hour treatment with MG132 at concentrations ranging from 5 to 10 µM is capable of accumulating cells in healthy metaphase. Depending on the desired accumulation of anaphase and telophase cells, either 5 µM or 10 µM MG132-treated cells can be utilized, tailored to the specific timing requirements of subsequent experiments. While 5 µM MG132-treated cells exhibit a higher rate of proper anaphase progression compared to those treated with 10 µM MG132 upon washout, these cells promptly progress into anaphase upon removal of the compound. On the other hand, 10 µM MG132-treated cells offer slightly more time for the preparation of subsequent procedures.

### Limitation of this study

For our synchronization method, we optimized the protocol using the RPE1 cell line, a normal, non-transformed human cell line expressing wild-type p53 (Bowden *et al*, 2020). It has been reported that certain inhibitors, particularly Cdk inhibitors, exhibit varying efficacies across different cell lines (Johnson *et al*., 2021; Trotter & Hagan, 2020). This variability may be attributed to the differential activities of Cdks in distinct cell types. A study demonstrated that in cancer cells, Cdk2 can compensate for the loss of Cdk1 during mitotic entry when Cdk1 is rapidly degraded using the auxin-degron system (Lau *et al*., 2021). However, this compensation does not occur in normal cells. While our optimized inhibitor concentrations provide a useful reference, adjustments may be required when applied to other cell lines.

## Discussion

Cell cycle synchronization is a commonly used method to accumulate cell populations in specific stages of the cell cycle to study stage-specific mechanisms and regulations. To achieve this, treatments with inhibitors targeting cell cycle-specific and essential kinases or proteins are commonly used (Dickson & Schwartz, 2009; Mills *et al*., 2017). However, these inhibitors often induce irreversible effects at higher concentrations and demonstrate inefficacy at lower concentrations. To precisely study cell cycle-specific mechanisms, it is critical to concentrate cells in the target cell cycle stage under conditions that are both healthy and reversible. Most characterizations of these inhibitors were performed using flow cytometry-based assays. Combining our immunofluorescence-based cell cycle identification method with cell synchronization (and washout), we demonstrate that all inhibitors we have tested induced certain defects and resulted in irreversibly arrested cells in reproductive cycles (**Fig. 1-5**). It is critical to minimize these effects for further experiments and quantification by using appropriate concentrations. For example, RPE1 cells synchronized in the G1 phase using optimal concentrations of Palbociclib still exhibited 20-30% arrested cells in the G1 phase 18 hours after washout (**Fig. 1c**). Similarly, cells synchronized to the G1 phase by Aphidicolin also exhibited ∼20% cells in the G1 phase 6 hours after washout (**Fig. 2b and 2c**). Surprisingly, RO-3306 is now more frequently used for G2 synchronization. Higher than optimal concentrations showed no G2 phase synchronization (**Fig. 3c**), indicating that high concentrations of RO-3306 might inhibit other Cdks, although RO-3306 is considered a selective inhibitor for Cdk1 (Jorda *et al*., 2018). Cells treated with the optimal concentration of RO-3306 can significantly accumulate in the G2 phase (approximately 60%); however, only 50% of these G2 phase cells can immediately enter mitosis after washout (**Fig. 3e**). Treatment with MG132 for more than one hour causes irreversible defects in metaphase cells, both with and without washout (**Fig. 5**). We summarize our recommended conditions for cell synchronizations at each stage of the cell cycle in RPE1 cells in **Supplementary Table 1**.

We demonstrated that all the inhibitors we tested were unable to prevent irreversible effects or other defects. This may be due to off-target effects of the inhibitors or difficulties in achieving complete washout. To circumvent these issues, developing conditional knockout cell lines for cell cycle kinases could be a viable alternative, although it requires additional effort to generate these strains. Notably, a previous study demonstrated that rapid depletion of Cdk1 in HeLa cells still allowed entry into mitosis, as Cdk2 compensates for Cdk1’s role in mitotic entry but not mitotic exit (Lau *et al*., 2021). Interestingly, RO-3306 effectively arrested HeLa cells in the G2 phase (Vassilev *et al*., 2006). This might be because RO-3306 inhibits not only Cdk1 but also other Cdks. This suggests that the use of inhibitors can effectively arrest cells at a specific point in the cell cycle, overcoming potential compensatory effects by other kinases. This approach may be more effective than using conditional knockout cell lines for targeting cell cycle kinases in certain cell types. Nevertheless, our detailed analysis of cell cycle inhibitors and the optimization of reversible and effective cell synchronization in RPE1 cells will provide a standard and serve as a reference for future research.

## Acknowledgement

We would like to thank Yu-Chia Chen, Yuhi Hara, Takanori Tsuchiya, and Tokai Hit for valuable suggestions, critical equipment and technical support. Part of this work is supported by Wisconsin Partnership Program, Research Forward from the Office of the Vice Chancellor for Research and Graduate Education (OVCRGE), start-up funding from University of Wisconsin-Madison SMPH, UW Carbone Cancer Center, and McArdle Laboratory for Cancer Research, and NIH grant R35GM147525 (to A.S.).

## Author contribution

YL.C. conducted precision imaging experiments and analyses, with assistance from S.R. and A.S. A.S. conceptualized, supervised, and funded the project. A.S. prepared the initial manuscript draft. All authors reviewed and contributed to the manuscript’s refinement.

## Methods

### Cell Culture

Human RPE1 cells were originally obtained from the American Type Culture Collection (ATCC, Manassas, VA, USA). RPE1 H2B-EGFP cells were obtained from Dr. Beth Weaver. RPE1 and RPE1 H2B-EGFP cells were grown in DMEM high glucose (Cytiva Hyclone; SH 30243.01) supplemented with 1% penicillin-streptomycin, 1% L-glutamine, and 10% fetal bovine serum under 5% CO2 at 37°C in an incubator.

### Cell Synchronization

Cells were plated one day prior to inhibitor treatment, reaching 60-70% confluency during the logarithmic growth phase at the time of treatment. Inhibitors used for cell cycle synchronization included Palbociclib, Aphidicolin, RO-3306, STLC, and MG132, detailed in **Supplementary Table 1**. Specifically, cells were synchronized at the G1 phase by incubating with Palbociclib for 24 hours. For S phase synchronization, cells were treated with Aphidicolin for 24 hours, followed by a washout, with collections at 4 or 6 hours post-washout. G2 phase synchronization involved a 24-hour incubation with RO-3306. For synchronization at metaphase, anaphase, and telophase, cells were treated with MG132 for 1 hour following a 24-hour RO-3306 treatment.

### Live-cell imaging

RPE1 H2B-EGFP cells were plated on 4-chamber 35mm glass bottom dishes (4 chamber with #1.5 glass, Cellvis) or µ-slide 8 well high glass bottom (ibidi, 80807) at least one day prior to imaging. After 24 hours of plating, cells were treated with inhibitors for cell synchronization (see Cell synchronization section) and, if necessary, subjected to washout before commencing live- cell imaging. Live-cell imaging was performed using a Nikon Ti2 inverted microscope equipped with a Hamamatsu Flash v2 camera, spectra-X LED light source (Lumencor), Shiraito PureBox with a STXG stage top incubator (TokaiHit), and a Plan Apo 20x objective (NA = 0.75) controlled by Nikon Elements software. Cells were recorded at 37°C with 5% CO2 in a stage-top incubator using the feedback control function to accurately maintain temperature of growth medium (Tokai Hit, STXG model). For non-wash out conditions, images were recorded for ∼24 hours at 30 minutes intervals with three z-stack images acquired at steps of 3 μm for each time point. For washout experiments, most of images were recorded for 12-24 hours at 3 or 6 minutes intervals.

### Immunofluorescence

Accurate identification of cell cycle stages was achieved using ImmunoCellCycle-ID, a tool we recently developed (Chen *et al*., 2024). The following primary and secondary antibodies, along with a DNA dye, were utilized: anti-CENP-F (kindly gifted by Dr. Stephen Taylor), PCNA (Santacruz, sc-56), CENP-C (MBL, PD-030), DAPI (Sigma, D9542), Guinea Pig IgG-Alexa 647 (JacksonImmuno, 706-606-148), Sheep IgG-Rhodamine Red X (JacksonImmuno, 713-546-147), and Mouse IgG (JacksonImmuno, 715-546-150). RPE1 cells were fixed by 4% PFA (Sigma) or 100% Methanol. Cells which fixed with PFA were then permeabilized by 0.5% NP40 (Sigma) and incubated with 0.1% BSA (Sigma). Stained samples were imaged by CSU W1 SoRa spinning disc confocal, which was equipped with Uniformizer and a Nikon Ti2 inverted microscope with a Hamamatsu Flash V2 camera and a 100x Oil objective (NA = 1.40). Microscope system was controlled by Nikon Elements software (Nikon).

### Mitotic shake-off

RPE1 cells were treated with 5 µM of STLC for 24 hours, after which mitotic cells were collected by shaking. The growth medium was then centrifuged to concentrate the cells. Subsequently, these cells were cytospin onto coverslips, fixed with 4% PFA, and stained with DAPI (refer to the **Immunofluorescence section** for details).

### Image analysis

Image analysis was performed using Nikon Elements software (Nikon) or Metamorph (Molecular Devices).

### Statistics

All experiments were independently repeated 2-3 times for mitotic duration measurements. p-values were calculated using one-way ANOVA and the two-tailed Student’s t-test. p-values < 0.05 were considered significant.

